# PEONY: a global reference database of DNA viral (vOTU) sequences from viral size-fractionated metagenomes (viromes)

**DOI:** 10.64898/2026.07.09.737585

**Authors:** HM Goemann, L Stern, M Perry, LS Hillary, JB Emerson

## Abstract

We present an update to the PIGEON (Phages and Integrated Genomes Encapsidated or Not) reference database of DNA viral sequences (vOTUs, mostly dsDNA bacteriophages) from global ecosystems. To reflect the inclusion of only virus size-fractionated metagenome- (virome-)derived vOTUs, we reintroduce the database as PEONY (Phages Encapsidated ONlY) and present new data summarizing the utility of this database.

## Introduction

Knowledge of the global biogeography of DNA viruses, particularly dsDNA bacteriophages (phages, viruses of bacteria) is rapidly expanding. Opportunities to synthesize these efforts are critical to understanding fundamental viral ecology, phage-host dynamics, and links to global biogeochemical cycling. Existing reference datasets focused on phages^1–5^ comprise a mix of viral sequences predicted from total DNA metagenomes and RNA metatranscriptomes, isolated viral genomes, and viral size-fractionated metagenomes (viromes). Ma et al.^1^ acknowledged that viromics is likely the most appropriate method for viral biogeography, as virions are separated from larger microbes to significantly increase viral recovery^6^. However, at the time of that publication, there were too few viromes available for exploring large-scale patterns in their Global Soil Virome dataset^1^. Here we contribute to this gap in publicly available viral databases by updating PIGEON, which is among the largest dsDNA viromics-specific databases to date.

PIGEON1.0^7^ was first assembled and made publicly available in 2021 with 266,125 viral operational taxonomic units (vOTUs > 10 kbp, clustered at 95% average nucleotide identity, ANI)^8^ from peat, soil, freshwater, marine, mammalian digestive, and plant-associated environments. The database was updated in 2023 (PIGEON2.0)^9^ to include additional vOTUs totaling 515,763. Due to the limited numbers of sequenced viromes at the time, particularly from soil, PIGEON1.0 and 2.0 included vOTUs recovered from both viromes and total DNA metagenomes.

## Methods

To complete this update, we compiled new virome datasets from our own recently published and as-yet unpublished work and from a Web of Science literature search. Inclusion/exclusion criteria for new studies are detailed in the Supplemental Information on the database Github repository. Briefly, we required assembled contigs or vOTUs generated from viral size-fractionated metagenomes (< 0.2 µm or < 0.45 µm). While there were additional studies that met our laboratory methodological inclusion criteria, we were unable to support large-scale re-assembly from raw data. Viral contigs and vOTUs were re-processed for viral sequence prediction via geNomad v1.9.4^10^ for consistency and filtered to > 10,000 bp length. We then individually (per-study) dereplicated, aligned, and clustered the contigs into vOTUs with vclust v1.3.1^11^ (leiden algorithm, ANI threshold 0.95, coverage threshold 0.85)^8^ prior to clustering with PIGEON2.0 to generate PIGEON3.0 (1,147,011 vOTUs). After removing 253,851 vOTUs from total metagenomes in PIGEON2.0 and reclustering with new the data, 945,661 vOTUs remained in PEONY.

## Results

PEONY vOTUs encompass terrestrial, aquatic, and mammalian digestive ecosystems across five continents (Fig. 1). Notably, 65% of vOTUs in PEONY originate from California, USA, where soils from various ecosystems (agricultural fields, grasslands, forests, wetlands, chaparral, and plant rhizospheres) have been routinely sampled by the Emerson Lab at the University of California, Davis and collaborators in recent years. This database provides a useful tool for studies of dsDNA viral (presumably, mostly bacteriophage) diversity and environmental distributions. Researchers can use PEONY to compare viruses from individual studies to global distribution patterns.

**Figure 1.**
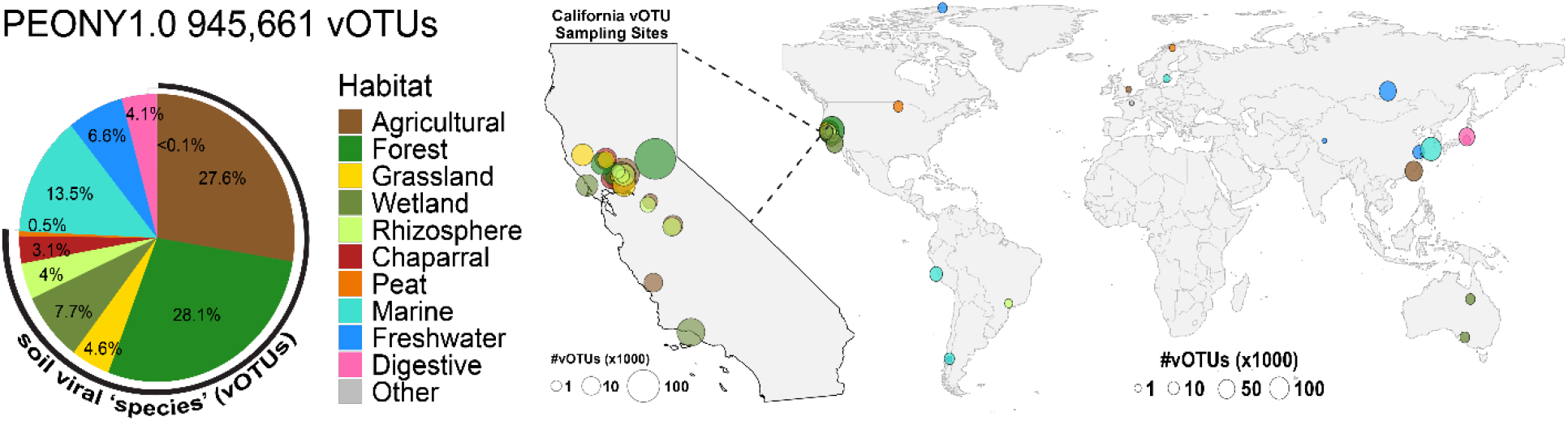
Distribution of PEONY vOTUs across ecosystems, both within California, USA, and globally. Size of mapped dots indicates the number of vOTUs (x1000) recovered from samples originating at a given location.

The code for database visualization in R, scripts for curation, and additional methods information are available on GitHub (https://github.com/hgoem/PEONY), and the databases (both PIGEON3.0 and PEONY) and associated metadata are available on Dryad (https://datadryad.org/10.5061/dryad.prr4xgz35).

## Notes

https://datadryad.org/dataset/doi:10.5061/dryad.prr4xgz35

https://github.com/hgoem/PEONY

### Competing Interest Statement

The authors have declared no competing interest.

