## Supplemental Information for "PEONY: a global reference database of DNA viral (vOTU) sequences from viral size-fractionated metagenomes (viromes)"

Running title: PEONY database

Authors: Goemann HM^1^, Stern L^1^, Perry M^1^, Hillary LS^1,2^, Emerson JB^1*^

Affiliations:

1. Department of Plant Pathology, University of California, Davis, Davis, CA, USA

2. School of Environmental & Natural Sciences, Bangor University, Bangor, Gwynedd, UK

Keywords:

Bacteriophage, biodiversity, viromics, size-fractionated metagenomics, database

Supplementary Information:

Literary Search

A *Web of Science* search was created on 12/01/25, using the below search terms, which yielded 3109 studies. The studies were split equally and randomly amongst four researchers (the first four authors of this resource announcement), using the R^1^ metagear package^2^ for Y/N/M (yes/no/maybe) screening for potential inclusion. The studies underwent a preliminary, manual title-abstract screening to eliminate total metagenomes, targeted sequencing, reviews, perspectives, spike-ins, and RNA-exclusive studies, resulting in 689 studies for full screening.

*Search terms:*

*TOPIC: Virom* OR “viral metagenom*” OR metavirome AND (“viral fraction” OR "0.22 µm" OR “0.2 µm” OR "0.45 µm" OR "0.1 µm" OR ultracentrifugation OR “iron chloride” OR ultracentrifuge OR TFF OR “tangential flow” OR PEG)*

*Date period: 2021-present*

The remaining 689 studies were re-randomly distributed amongst the same four researchers so that every manuscript was screened by two different researchers between the first and second screening efforts. The inclusion criteria are listed below:

Studies were included if samples were size-fractionated viral metagenomes (< 0.22 µm or < 0.45 µm) and readily available at the assembled contig or clustered vOTU level. No reviews, perspectives, meta-analyses, mining from total metagenomes, exclusively RNA studies, multiple-displacement amplification (MDA), or dual RNA/DNA extractions followed by reverse transcription and combined sequencing were included. This screening resulted in a list of 16 studies^3–19^ that underwent bioinformatic processing. For datasets from which the data availability was unclear, the corresponding authors were emailed with a request for data access, which resulted in the inclusion of two additional studies^19,20^. The full list of search results and subsequent screening are included on the project Github.

Inclusion of new and as-yet unpublished data from the Emerson Lab:

All published studies from the Emerson lab that met the laboratory and sample processing inclusion criteria above were included. In addition, unpublished viromic datasets with presumed final vOTU sequences were included, at the discretion of both Emerson and the current first author of each study. Further information is reported in the supplementary table on Github.
